# The ADP-glucose pyrophosphorylase from *Melainabacteria*: a comparative study between photosynthetic and non-photosynthetic bacterial sources

**DOI:** 10.1101/2021.06.18.448987

**Authors:** María V. Ferretti, Rania A. Hussien, Miguel A. Ballicora, Alberto A. Iglesias, Carlos M. Figueroa, Matías D. Asencion Diez

## Abstract

Until recently, all members of the cyanobacterial phylum were considered capable of performing oxygenic photosynthesis. This view has been questioned after the discovery of a group of presumed non-photosynthetic cyanobacteria named *Melainabacteria*. Using metagenomic data, we identified sequences encoding putative ADP-glucose pyrophosphorylase (EC 2.7.7.27, ADP-GlcPPase) from free-living and intestinal *Melainabacteria*. These genes were *de novo* synthesized and overexpressed in *Escherichia coli*. The purified recombinant proteins from the free-living and the intestinal *Melainabacteria* showed ADP-GlcPPase activity, with *V*_max_ values of 2.3 and 7.1 U/mg, respectively. Both enzymes had similar affinities towards ATP (*S*_0.5_ ∼0.3 mM) although the one from the intestinal source displayed a 6-fold higher affinity for glucose-1P. Both recombinant ADP-GlcPPases were sensitive to allosteric activation by glucose-6P (*A*_0.5_ ∼0.3 mM), and to inhibition by Pi and ADP (*I*_0.5_ between 0.2 to 3 mM). Interestingly, the enzymes from *Melainabacteria* were insensitive to 3-phosphoglycerate, which is the principal activator of ADP-GlcPPases from photosynthetic cyanobacteria. To the best of our knowledge, this is the first biochemical characterization of an active enzyme from *Melainabacteria*, offering further data to discussions regarding their phylogenetic position. This work contributes to a better understanding regarding the evolution of allosteric mechanisms in ADP-GlcPPases, an essential enzyme for the synthesis of glycogen in prokaryotes and starch in plants.

## Introduction

Most living organisms produce α-1,4-glucans as a strategy to store carbon and energy, which can be mobilized under conditions of nutrient deficiency. Bacteria and heterotrophic eukaryotes accumulate glycogen, whereas starch is the reserve carbohydrate in green algae and higher plants [1]. The build-up of glycogen and starch in bacteria and plants, respectively, involves a similar pathway where ADP-glucose (ADP-Glc) is the glycosyl donor for polysaccharide elongation. In such metabolic route, the sugar nucleotide synthesis is the rate-limiting step, catalyzed by ADP-Glc pyrophosphorylase (ADP-GlcPPase, EC 2.7.7.27). Indeed, ADP-GlcPPase is allosterically regulated by metabolites from the central carbon utilization pathway in the respective organism [1–4]. In this context, the enzyme from *Escherichia coli* is mainly activated by fructose-1,6-bisphosphate, a key intermediate in the Embden-Meyerhof route. At the same time, fructose-6-phosphate (Fru-6P) and pyruvate are the principal activators of the enzyme from *Agrobacterium tumefaciens*, where the Entner-Doudoroff is the main glycolytic pathway [5–9]. Similarly, ADP-GlcPPases from organisms performing oxygenic photosynthesis (cyanobacteria, green algae, and higher plants) are primarily activated by 3-phosphoglycerate (3-PGA) and inhibited by inorganic orthophosphate (Pi) [1,10–12].

Cyanobacteria are a highly diverse group of Gram-negative prokaryotes that colonized a wide range of environments, from desert crusts to fresh and marine waters and from the tropics to the poles [13]. These microorganisms modified the Earth’s atmosphere through oxygenic photosynthesis, which enabled the evolution of life into more complex forms [14]. Photosynthetic cyanobacteria have been studied for decades, and their diversity is described in terms of both morphology and genetics [15–17]. With the development of metagenomics, an unexpected variety of organisms was unveiled in many ecosystems, including non-photosynthetic bacteria closely related to the clade Cyanobacteria [18–21].

One group of these microorganisms was named *Melainabacteria* [19] because several representatives of this cluster were found in aphotic environments. Firstly, they were considered as a sister phylum of Cyanobacteria [19]; later, data from genomic sequences analysis suggested that *Melainabacteria* belongs to the superphylum of Cyanobacteria [20], although these suggestions have not been validated so far. Based on this evidence, a new classification has been proposed for the phylum Cyanobacteria. This arrangement includes the class-level lineages *Oxyphotobacteria* (cyanobacteria performing oxygenic photosynthesis) and *Melainabacteria*, as well as a third class called ML635J-21 [20], recently named *Sericytochromatia* [22]. After diverging from *Melainabacteria*, the *Oxyphotobacteria* developed oxygenic photosynthesis around 2.4–2.35 billion years ago, as estimated from the molecular clock and geological data [22–24].

*Melainabacteria*, described for the first time only a few years ago, is a group of poorly characterized anaerobic bacteria. To date, only a handful of *Melainabacteria* genomes have been sequenced [22], and thus, the metabolism, biological functions, and ecological roles of these organisms are not fully known. Representatives of *Melainabacteria* have been found in photic and aphotic environments such as (*i*) sub-surface groundwater [19], (*ii*) lake water and algal biofilms [22,25], (*iii*) marine and lacustrine sediment [26], and (*iv*) animal and human faeces [20] and guts [26]. All sequenced genomes of *Melainabacteria* confirm they lack the entire photosynthetic apparatus, which may support the hypothesis that acquisition of photosystems in *Oxyphotobacteria* occurred after divergence from the non-photosynthetic *Melainabacteria* [20,22]. Consequently, the characterization of enzymes from main metabolic segments (such as the synthesis of the energy/carbon storage molecule glycogen) is critical to sum biochemical criteria to help the scientific community to further classify this group of bacteria.

By studying the biochemical properties of cyanobacterial ADP-GlcPPases, an evolutionary thread could be established between bacterial glycogen and starch synthesis metabolism [1,2,27,28]. Hence, the particular regulatory properties of ADP-GlcPPases prompted us to explore the features of this enzyme in *Melainabacteria*. In this framework, we *de novo* synthesized the genes encoding ADP-GlcPPases from intestinal (in*Mel*GlgC) and free-living (fl*Mel*GlgC) *Melainabacteria*. The recombinant proteins were produced, purified, and kinetically characterized. For the sake of comparison, we also made the homologous enzyme from photosynthetic *Anabaena* PCC 7120 (*Ana*GlgC). Our results indicate that *Melainabacteria* ADP-GlcPPases have distinctive kinetic and regulatory properties, which might fit a heterotrophic metabolism commonly found in diverse bacterial organisms.

## Material and Methods

### Chemicals, bacterial strains and plasmids

Chemicals used for enzymatic assays were from Sigma-Aldrich (St. Louis, MO, USA). All the other reagents were of the highest quality available. *Escherichia coli* Top 10 (Invitrogen) were used for plasmid maintenance. The *glgC* genes from *Anabaena* PCC 7120, intestinal and free-living *Melainabacteria* were expressed in *E. coli* BL21 (DE3) (Invitrogen) using the pET28b vector (Novagen). DNA manipulations, molecular biology techniques, and *E. coli* cultivation and transformation were performed according to standard protocols [29].

### Phylogenetic analysis

Amino acid sequences of ADP-GlcPPases from different organisms were downloaded from the NCBI database (http://www.ncbi.nlm.nih.gov) and classified into different groups using taxonomic data provided by the NCBI. Sequences were manually curated to remove duplicates and near-duplicates (i.e., mutants and strains from the same species). We constructed a preliminary alignment using the ClustalW multiple sequence alignment server [30], which was then manually refined with the BioEdit 7.0 program [31]. Tree reconstruction was performed using the neighbour-joining algorithm with a bootstrap of 1,000 in the program SeaView 4.3 [32]. The phylogenetic tree was prepared with the FigTree 1.3 program (http://tree.bio.ed.ac.uk/software/figtree/).

### *Cloning of* glgC *genes*

Genes encoding ADP-GlcPPases from intestinal and free-living *Melainabacteria* were *de novo* synthesized (Bio Basic, Canada) according to genomic information for these bacteria [19,21], available in the NCBI database (Nucleotide IDs CP017245.1 and MFRL00000000.1, respectively). The genes encoding intestinal and free-living *Melainabacteria* GlgC proteins (NCBI Protein IDs AOR37842.1 and OGI00355.1, respectively) were optimized for expression in *E. coli* and inserted into the pET28b vector between the *Nde*I and *Sac*I restriction sites, to produce the recombinant proteins with an N-terminal His-tag. The same procedure was performed for the gene encoding the *Anabaena* PCC 7120 GlgC protein (NCBI Protein ID WP_010998776.1).

### Enzyme production and purification

Transformed *E. coli* BL21 (DE3) were grown in YT2X medium (16 g/l tryptone; 10 g/l yeast extract; 5 g/l NaCl) supplemented with kanamycin (50 µg/ml) at 37 °C and 200 rpm, until reaching an optical density at 600 nm of ∼0.6. Recombinant protein expression was induced with 0.1 mM isopropyl-β-D-1-thiogalactopyranoside for 16 h at 18 °C. Cells were harvested by centrifugation at 5000 × *g* for 10 min and stored at -20 °C until use.

His-tagged proteins were purified at 4 °C by immobilized metal affinity chromatography (IMAC). Cells were resuspended in *Buffer H* [50 mM Tris-HCl pH 8.0, 300 mM NaCl, 10mM imidazole, 5% (v/v) glycerol] and disrupted by sonication. The suspension was centrifuged twice at 30000 × *g* for 10 min, and the supernatant (crude extract) loaded on a 1-ml His-Trap column (GE Healthcare) previously equilibrated with *Buffer H*. The recombinant proteins were eluted with a linear gradient from 10 to 300 mM imidazole in *Buffer H*. Fractions containing the highest activity were pooled, concentrated to 2 ml, and dialyzed against *Buffer S* [50 mM HEPES-NaOH, 10% (w/v) sucrose, 0.2 mM DTT, 1 mM EDTA]. The resulting enzyme samples preparations were stored at -80 °C until use, remaining fully active for at least 10 months.

### Protein methods

Protein concentration was determined with the Bradford reagent [33], using bovine serum albumin (BSA) as a standard. The purity of the recombinant proteins was assessed by sodium dodecyl sulfate-polyacrylamide gel electrophoresis (SDS-PAGE), according to Laemmli [34]. Gels were loaded with 5 to 50 µg of protein per well and stained with Coomassie Brilliant Blue.

### Native molecular mass determination

The native molecular mass of the recombinant proteins was determined by gel filtration using a Superdex 200 10/300 column (GE Healthcare), previously calibrated with protein standards (GE Healthcare), including thyroglobulin (669 kDa), ferritin (440 kDa), aldolase (158 kDa), conalbumin (75 kDa), and ovalbumin (44 kDa). The void volume of the column was determined using Dextran Blue (Promega).

### Enzyme activity assays

ADP-GlcPPase activity was determined at 37 °C in the direction of ADP-Glc synthesis, following the formation of P_i_ after hydrolysis of PP_i_ by inorganic pyrophosphatase, using a highly sensitive colorimetric method [35]. Reaction mixtures contained (unless otherwise specified) 50 mM MOPS-NaOH pH 8.0, 10 mM MgCl_2_, 1.5 mM ATP, 0.2 mg/ml BSA, 0.5 U/ml yeast inorganic pyrophosphatase and a proper enzyme dilution. Assays were initiated by the addition of 1.5 mM Glc-1P in a total volume of 50 µl. Reaction mixtures were incubated for 10 min at 37 °C and terminated by adding 400 µl of the Malachite Green reagent. The complex formed with the released P_i_ was measured at 630 nm in a 96-well microplate reader (Multiskan GO, Thermo).

To test P_i_ inhibition, ADP-GlcPPase activity was measured using a coupled-enzyme spectrophotometric assay. Reaction mixtures contained 50 mM MOPS-NaOH pH 8.0, 10 mM MgCl_2_, 0.3 mM phospho*enol*pyruvate, 0.3 mM NADH, 2 mM ATP, 1 mg/ml rabbit muscle glycogen, 0.8 U/µl *E. coli* glycogen synthase, 0.1 U/µl pyruvate kinase, 0.02 U/µl lactate dehydrogenase, 0.2 mg/ml BSA and enzyme in a total volume of 50 μl. The reaction was initiated with 2 mM Glc-1P, and activity was measured by following NADH oxidation at 340 nm and 37 °C using a 384-microplate reader (Multiskan GO, Thermo). One unit of activity (U) is defined as the amount of enzyme catalyzing the formation of 1 µmol of product per min, under the above specified conditions.

Saturation curves were constructed by assaying enzyme activity at different concentrations of the variable substrate or effector, while the others remained at saturating levels. Plots of enzyme activity (U/mg) *versus* substrate (or effector) concentration (mM) were used to calculate the kinetic constants, by fitting the experimental data to a modified Hill equation [36]. Fitting was performed with the Levenberg-Marquardt non-linear least-squares algorithm provided by the computer program Origin 8.0 (OriginLab). Accordingly, we calculated the Hill coefficient (*n*_H_), the maximal velocity (*V*_max_), and the concentrations of activator, substrate or inhibitor giving 50% of the maximal activation (*A*_0.5_), velocity (*S*_0.5_) or inhibition (*I*_0.5_), respectively. All kinetic constants are the mean of at least three independent sets of data, which were reproducible within a range of ± 10%.

## Results

### *Identification of* glgC *genes, molecular cloning, and phylogenetic analysis of ADP-GlcPPases from* Melainabacteria

We identified *glgC* genes, encoding putative ADP-GlcPPases, in metagenomic databases from intestinal (*inMelglgC*) and free-living (*flMelglgC*) *Melainabacteria* [19,21]. The *inMelglgC* (1,236 bp) and *flMelglgC* (1,233 bp) genes code for proteins of ∼45 kDa, which share identities of 66.18% between them; ∼35% with *Anabaena* GlgC; ∼43% with the *A. tumefaciens* GlgC; and ∼44% with the *S. coelicolor* GlgC. Further to this comparison, it is worth considering the already established structure to function relationships between ADP-GlcPPases regarding central metabolism in one organism [1,2]. In this context, we extended the comparison of the GlgCs’ amino acid sequences, obtaining the phylogenetic tree shown in Figure 1. As shown, the analysis protein found GlgCs from *Melainabacteria* in a cluster separated from those present in bacteria performing oxygenic photosynthesis. Indeed, *Melainabacteria* ADP-GlcPPases locate closer to proteins from heterotrophic bacteria, particularly Actinobacteria. These results trigger additional biological and evolutionary questions since the taxonomic classification of *Melainabacteria* [20], and the phylogenetic position of their ADP-GlcPPases are markedly different (Figure 1).

**Figure 1.**
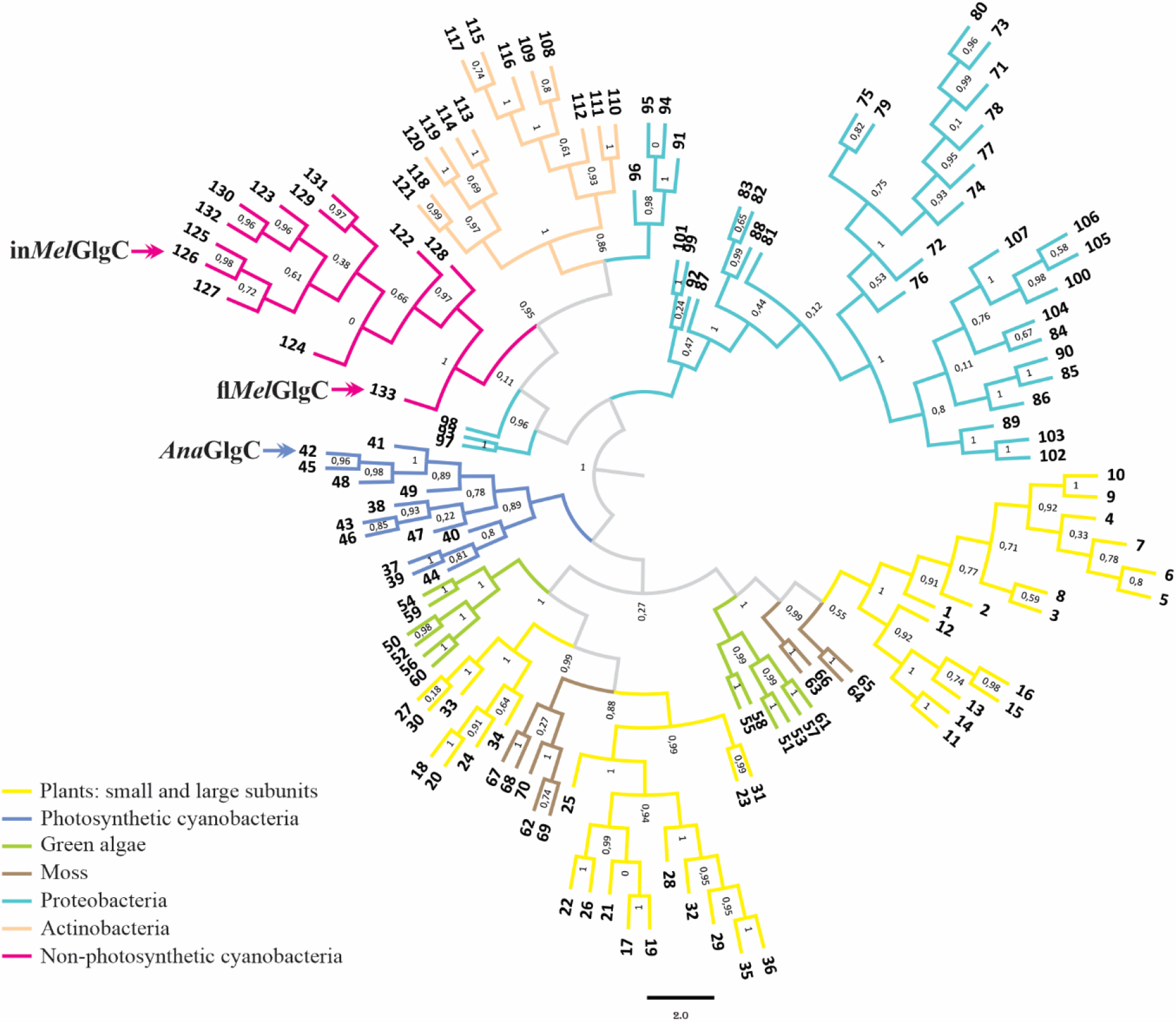
Phylogenetic tree of GlgC from different organisms. The tree was built as described in Materials and Methods. Protein sequences are numbered with codes indexed in Supplemental Table S3.

To explore beyond the phylogenetic relationships between ADP-GlcPPases, we *de novo* synthesized the genes in*MelglgC* and fl*MelglgC* to produce and characterize the respective recombinant proteins. For comparison, we followed the same approach to obtain the enzyme from *Anabaena* PCC 7120, which was already characterized in detail [11]. Supplemental Figure 1A illustrates that the ADP-GlcPPases from both *Melainabacteria* (in*Mel*GlgC and fl*Mel*GlgC), as well as that from *Anabaena* PCC 7120 (*Ana*GlgC), were obtained with a high purity level. The purified enzymes exhibited specific activity values of 7.1 (in*Mel*GlgC), 2.3 (fl*Mel*GlgC), and 0.31 (*Ana*GlgC) U/mg (in the absence of allosteric activators). Both ADP-GlcPPases from *Melainabacteria* eluted from the gel filtration column with molecular masses between 180 and 190 kDa (Supplemental Figure 1B). Considering the theoretical mass of these proteins and results obtained by SDS-PAGE (Supplemental Figure 1A), we conclude that both enzymes are homotetramers, which agrees with the quaternary structure of ADP-GlcPPases characterized so far [1,2].

### *Kinetic and regulatory properties of* Melainabacteria *ADP-GlcPPases*

The recombinant ADP-GlcPPases were kinetically characterized in the direction of ADP-Glc synthesis. Saturation curves for Glc-1P and ATP of the *Melainabacteria* enzymes showed deviation from the hyperbolic behavior (Supplemental Figure S2), with similar affinities towards ATP in both cases (Table 1). However, in*Mel*GlgC displayed a 6-fold lower *S*_0.5_ for Glc-1P than fl*Mel*GlgC (Table 1). In the comparative analysis, *Ana*GlgC exhibited an apparent affinity for Glc-1P one order of magnitude higher than fl*Mel*GlgC (Table 1). We also explored the effect of different metabolites, known to activate or inhibit ADP-GlcPPases from various organisms [1,2,9,37–39], on the activity of the *Melainabacteria* enzymes. As detailed in Figure 2, many of the assayed compounds exerted changes on the kinetics of in*Mel*GlgC and fl*Mel*GlgC. Among these, glucose-6P (Glc-6P), Fru-6P, and mannose-6P (Man-6P) activated, while ADP and Pi inhibited both enzymes (Figure 2). Noteworthy, the activity of *Melainabacteria* enzymes did not significantly change in the presence of 3-PGA (Figure 2), the primary activator of ADP-GlcPPases from oxygenic photosynthetic organisms [1,2,11], including those from cyanobacterial sources characterized so far [28,40,41].

**Table 1.**
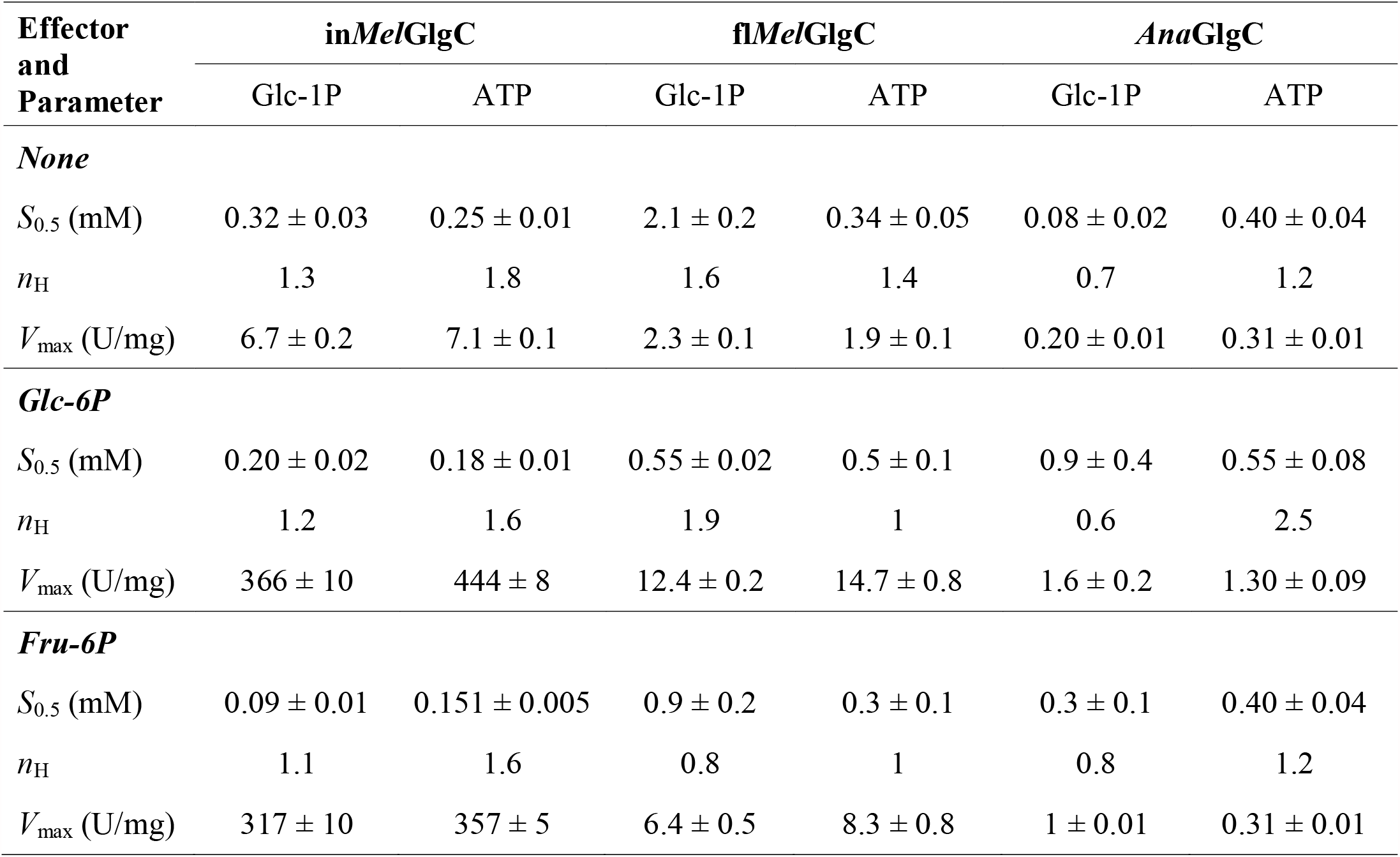
Kinetic parameters for the cyanobacterial ADP-GlcPPases characterized in this work in the absence and in the presence of allosteric activators. Activity was assayed as described in Materials and Methods. Kinetic parameters were calculated by the fitting software, using the mean of three independent datasets.

| Effector<br>and<br>Parameter | in*Mel*GlgC |  | fl*Mel*GlgC |  | *Ana*GlgC |  |
| --- | --- | --- | --- | --- | --- | --- |
|  | Glc-1P | ATP | Glc-1P | ATP | Glc-1P | ATP |
| ***None*** |  |  |  |  |  |  |
| $S_{0.5}$ (mM) | $0.32 \pm 0.03$ | $0.25 \pm 0.01$ | $2.1 \pm 0.2$ | $0.34 \pm 0.05$ | $0.08 \pm 0.02$ | $0.40 \pm 0.04$ |
| $n_H$ | 1.3 | 1.8 | 1.6 | 1.4 | 0.7 | 1.2 |
| $V_{\max}$ (U/mg) | $6.7 \pm 0.2$ | $7.1 \pm 0.1$ | $2.3 \pm 0.1$ | $1.9 \pm 0.1$ | $0.20 \pm 0.01$ | $0.31 \pm 0.01$ |
| ***Glc-6P*** |  |  |  |  |  |  |
| $S_{0.5}$ (mM) | $0.20 \pm 0.02$ | $0.18 \pm 0.01$ | $0.55 \pm 0.02$ | $0.5 \pm 0.1$ | $0.9 \pm 0.4$ | $0.55 \pm 0.08$ |
| $n_H$ | 1.2 | 1.6 | 1.9 | 1 | 0.6 | 2.5 |
| $V_{\max}$ (U/mg) | $366 \pm 10$ | $444 \pm 8$ | $12.4 \pm 0.2$ | $14.7 \pm 0.8$ | $1.6 \pm 0.2$ | $1.30 \pm 0.09$ |
| ***Fru-6P*** |  |  |  |  |  |  |
| $S_{0.5}$ (mM) | $0.09 \pm 0.01$ | $0.151 \pm 0.005$ | $0.9 \pm 0.2$ | $0.3 \pm 0.1$ | $0.3 \pm 0.1$ | $0.40 \pm 0.04$ |
| $n_H$ | 1.1 | 1.6 | 0.8 | 1 | 0.8 | 1.2 |
| $V_{\max}$ (U/mg) | $317 \pm 10$ | $357 \pm 5$ | $6.4 \pm 0.5$ | $8.3 \pm 0.8$ | $1 \pm 0.01$ | $0.31 \pm 0.01$ |

**Figure 2.**
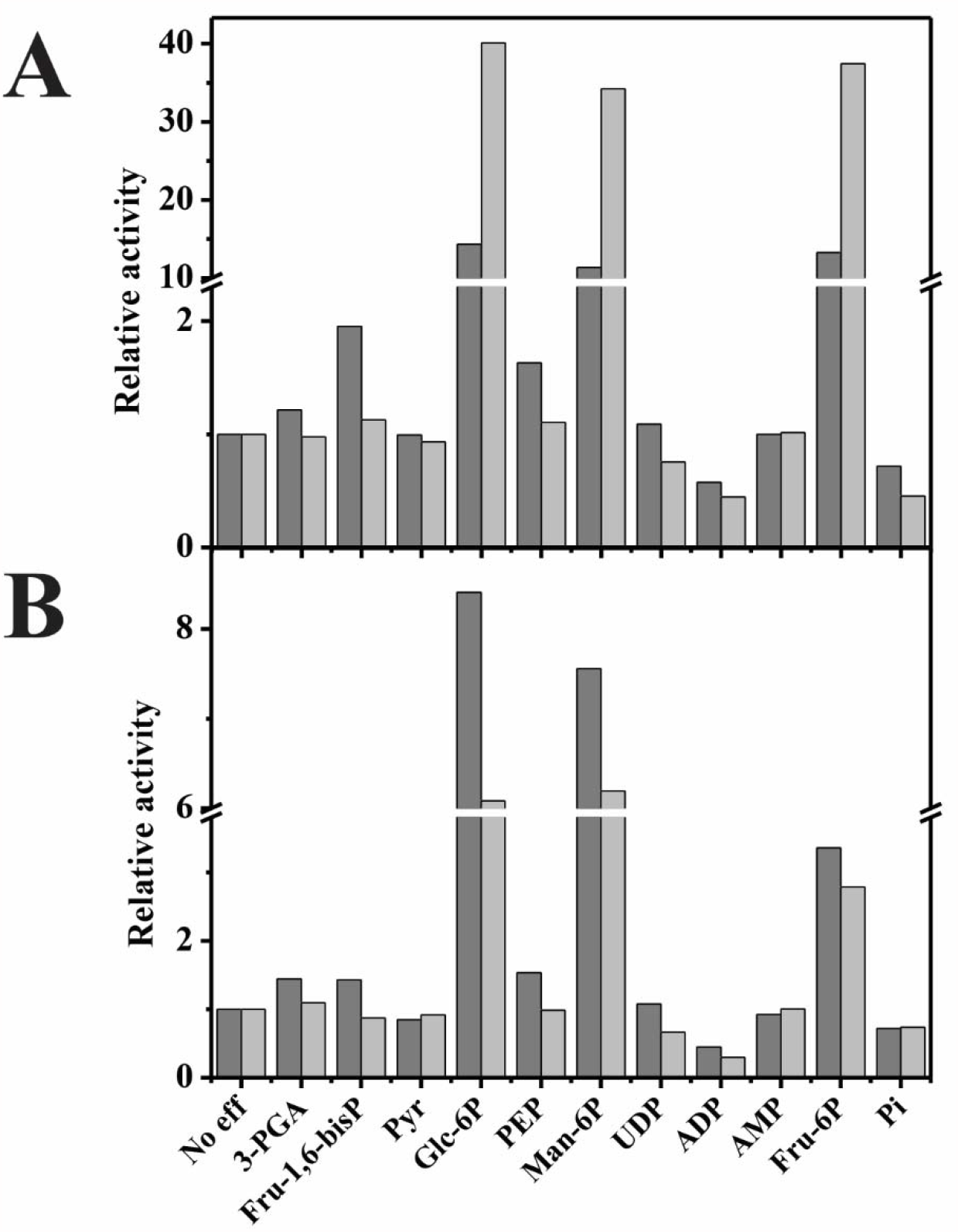
Effect of different metabolites on the activity of ADP-GlcPPases from non-photosynthetic cyanobacteria. **(A)** Intestinal *Melainabacteria* GlgC and **(B)** free-living *Melainabacteria* GlgC. Relative activities were calculated as the ratio between the activities in the presence and absence of the respective effector. The value of 1 corresponds to the respective ADP-GlcPPase *V*_max_ (see Table 1). The metabolite concentration was 2.5 mM in all cases. Assays were performed using two sets of Glc-1P and ATP concentrations: subsaturating (dark gray bars) or saturating (light gray bars).

We then performed a detailed study of the activation kinetics for the different ADP-GlcPPases studied in this work. As shown in Figure 3A, Glc-6P activated in*Mel*GlgC and fl*Mel*GlgC 54- and 12-fold, respectively, with similar *A*_0.5_ values (Supplemental Table S1). The activation of *Ana*GlgC by Glc-6P reached a maximum of 5-fold, and the relative affinity towards the hexose-P was 5-fold lower compared to the enzymes from *Melainabacteria* (Supplemental Table S1). Further analysis of substrate saturation kinetics showed that Glc-6P did not significantly alter the apparent affinities of the enzymes towards ATP. Instead, Glc-6P increased 1.6- and 3.8-fold the Glc-1P apparent affinities of in*Mel*GlgC and fl*Mel*GlgC, respectively; conversely, Glc-6P decreased 10-fold the Glc-1P apparent affinity in the case of *Ana*GlgC (Table 1). Fru-6P and Man-6P (respectively) activated in*Mel*GlgC (40- and 15-fold), fl*Mel*GlgC (4- and 13-fold), and *Ana*GlgC (15- and 6-fold), with *A*_0.5_ values in the range 0.5-1.5 mM (Supplemental Table S1). Figure 3B shows that 3-PGA has no effect on the activity of ADP-GlcPPases from *Melainabacteria* (up to 5 mM). At the same time, *Ana*GlgC was activated 30-fold (with an *A*_0.5_ of 0.2 mM), which agrees with previous work [11]. Inhibition kinetics confirmed that ADP and Pi are inhibitors of the studied enzymes (Supplemental Table S2), although Pi inhibition was more pronounced in *Ana*GlgC than in in*Mel*GlgC and fl*Mel*GlgC (*I*_0.5_ values were 0.09, 0.23, and 2.3 mM, respectively).

**Figure 3.**
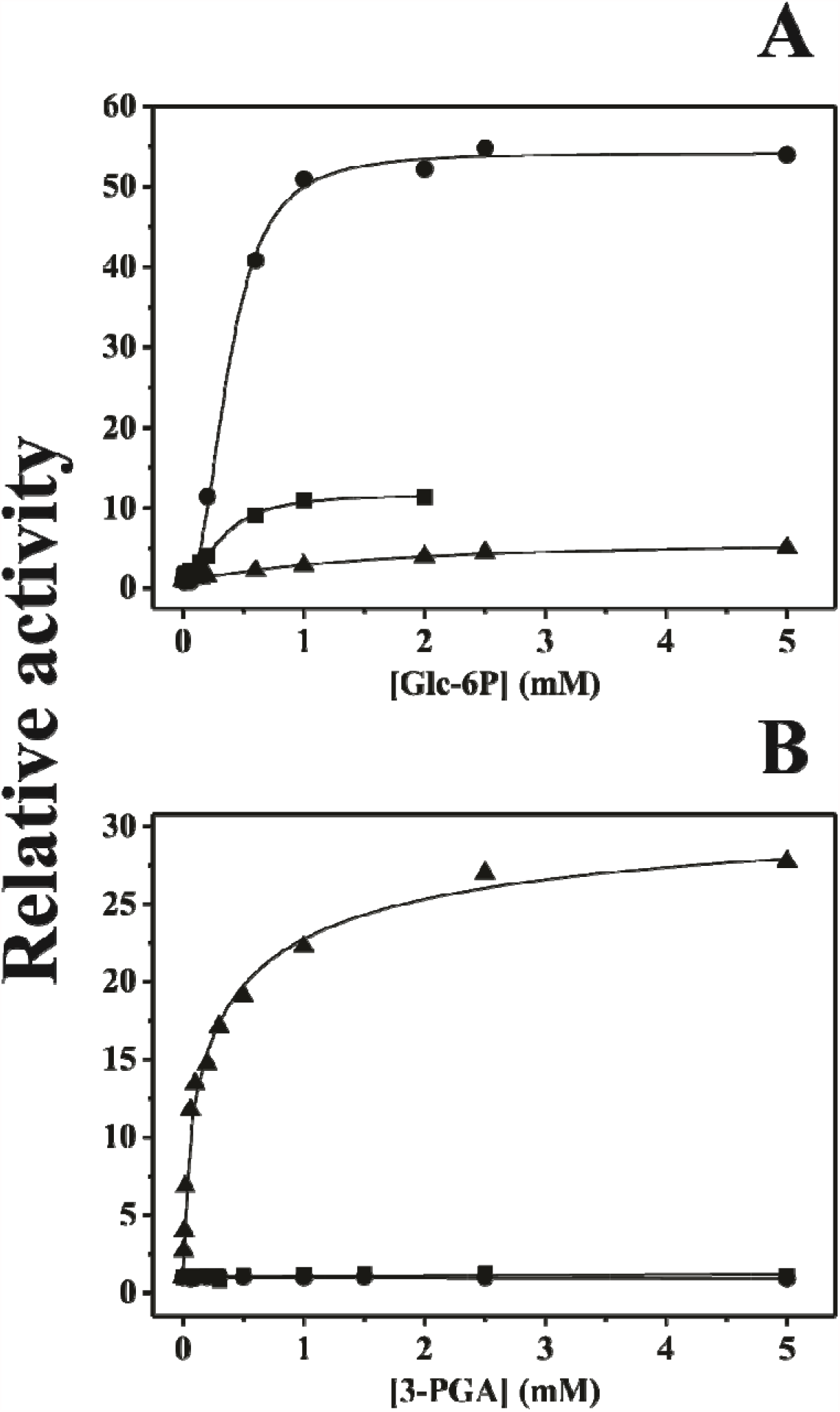
Activation of cyanobacterial ADP-GlcPPases by Glc-6P. **(A) and 3-PGA (B)**. Intestinal *Melainabacteria* GlgC (filled circles), free-living *Melainabacteria* GlgC (filled squares), *Anabaena* GlgC (filled triangle). The value of 1 corresponds to activities of 7.0, 2.0, and 0.25 U/mg for in*Mel*GlgC, fl*Mel*GlgC, and *Anabaena* ADP-GlcPPases, respectively.

## Discussion

It was initially thought that all members of the cyanobacterial phylum were capable of performing oxygenic photosynthesis [20,42]. This scenario was recently questioned after discovering *Melainabacteria*, a bacterial group closely related to Cyanobacteria at a phylogenetic level but incapable of performing photosynthesis [19,21]. The analysis of metagenomic information allowed us to find genes from *Melainabacteria* related to carbohydrate metabolism, particularly glycogen synthesis (see below and Table 2). Then, analyzing the kinetic and regulatory properties of key metabolic enzymes would contribute to the biochemical and evolutionary discussion concerning the classification of cyanobacteria and their sister-clades, such as *Melainabacteria*.

**Table 2.** Proteins detected after BLAST with data from *Melainabacteria*, compared to *Anabaena*.

| Gene | Intestinal<br>*Melainabacteria*<br>MELA1 | Free-living<br>*Melainabacteria*<br>GWF2_37_15 | *Anabaena*<br>PCC 7120 |
| --- | --- | --- | --- |
| *glgC* | ID: AOR37842.1 | ID: OGI00355.1 | ID: BAB76344.1 |
| *glgA* | ID: AOR37764.1 | ID: OGI01396.1 | ID: BAB73578.1<br>ID: BAB77555.1 |
| *glgB* | ID: AOR39099.1 | ID: OGI01242.1 | ID: BAB72670.1 |
| *glgX* | ID: AOR37731.1 | ID: OGI01270.1 | ID: BAB77692.1 |
| *glgP* | ID: AOR37860.1 | ID: OGI02531.1 | ID: BAB73229.1 |
| *glgE1* | ID: AOR39243.1 | ID: OGI01701.1<br>ID: OGI04937.1 | **not found** |
| *glgE2* | ID: AOR39243.1 | ID: OGI01701.1<br>ID: OGI04937.1 | **not found** |
| *sus* | **not found** | **not found** | ID: BAB76684.1 |
| *sps* | **not found** | **not found** | ID: BAB75069.1<br>ID: BAB72334.1 |

We focused on sequences encoding ADP□GlcPPase, which catalyzes the first committed step in the pathway for bacterial glycogen synthesis [2,4]. As previously mentioned, ADP-GlcPPases from Cyanobacteria performing oxygenic photosynthesis are activated by 3-PGA, while glycolytic intermediates control those from heterotrophic microorganisms, *e*.*g*., fructose-1,6-bisphosphate and pyruvate in the enzyme from *E. coli* [5–9]. Here, it is important to remark that ADP-GlcPPase’s regulatory properties are intimately related to main metabolic pathways in the organisms, then constituting a relevant issue to inferring on *Melainabacteria* metabolism. The comparative analysis of the structural, kinetic, and regulatory properties of different ADP-GlcPPases has contributed to understanding better the evolution of allosteric mechanisms in this family of biological catalysts [8,9,43–46]. In this regard, we sought to characterize ADP-GlcPPases from *Melainabacteria* since these bacteria seem to be located at a phylogenetic enclave which deserves further characterization. Hence, we constructed a phylogenetic tree to gain more information concerning the evolutionary relationship between *Melainabacteria* ADP-GlcPPases and enzymes from other taxonomic groups. Our phylogenetic analysis showed that *Melainabacteria* sequences were grouped closer to heterotrophic bacteria than to photosynthetic Cyanobacteria (Figure 1). ADP-GlcPPases from different prokaryotic sources showed to be homotetrameric, with subunits of about 45–50 kDa [2]. So far, the only exception is the enzyme from Firmicutes, a heterotetramer composed of two subunit types, GlgC and GlgD [2,47,48]. We could only find a single *glgC* gene in metagenomic data from *Melainabacteria* and, after recombinant expression of two different enzymes; we proved that both proteins are homotetramers (Supplemental Figure 1). Thus, ADP-GlcPPases from this Cyanobacteria sister-clade have a structural architecture similar to that from photosynthetic cyanobacteria (and most bacterial sources but Firmicutes), sustaining the importance of advancing with the kinetic and regulatory characterization of these enzymes to elucidate their structure-to-function relationships.

In a general view, the enzymes from *Melainabacteria* presented specific activities one order of magnitude higher than that from *Anabaena* PCC 7120 (in the absence of allosteric activators). However, the latter showed a higher apparent affinity towards ATP and Glc-1P (Table 1). Noteworthy, the catalytic capacity of ADP-GlcPPases from *Melainabacteria* was similar to that observed for enzymes from heterotrophic bacteria [1,2,49]. Interestingly, ADP-GlcPPases from *Melainabacteria* lack 3-PGA activation but are highly sensitive to hexose-6P (Glc-6P, Fru-6P, and Man-6P; Supplemental Table S1), similarly to the enzymes from heterotrophic bacteria [2], particularly those from Actinobacteria [37,39,50,51]. Remarkably, actinobacterial ADP-GlcPPases and those from *Melainabacteria* are located in the same branch of the phylogenetic tree (Figure 1). The activation by Glc-6P of in*Mel*GlgC (Figure 3) is the highest reported so far for this metabolite, while the effect on fl*Mel*GlgC is similar to that from the *Rhodococcus jostii* enzyme [50]. Given the proximity between ADP-GlcPPases from Actinobacteria and *Melainabacteria*, we foresee that these enzymes will be useful to explain the molecular mechanism underlying ADP-GlcPPase activation by Glc-6P. Even more, since Glc-6P is the common effector between ADP-GlcPPases from photosynthetic and some heterotrophic organisms (Figure 3 and Table 1), elucidating this allosteric mechanism will be critical to illuminate the evolutionary scenario related to changes on the sensitivity to a given effector.

Using the available metagenomic data, we also analyzed the existence in *Melainabacteria* of other genes related to glycogen metabolism in bacteria. As shown in Table 2, we found these organisms contain *glgA* and *glgB* genes, putatively encoding glycogen synthase (EC 2.4.1.21) and branching enzyme (EC 2.4.1.18). Curiously, the putative GlgA from *Melainabacteria* (so far, the only one) displays higher identity to the two homologous enzymes from *Synechocystis* PCC 6803, GlgA1 (37%) and GlgA2 (33%) - the latter probably related to glucan priming in some Cyanobacteria [52]-than to the one from *E. coli* (∼29%) or *Mycobacterium tuberculosis* (∼21%), recently renamed GlgM [53]. Besides, the GlgAs from *Melainabacteria* and *Anabaena* share a 36% identity between each other. Also, the *Anabaena* GlgA possess 73% and 30% identity with the GlgA1 and GlgA2 proteins from *Synechocystis*, respectively. Regarding the branching enzyme (GlgB), we found a protein sequence in *Melainabacteria* with a 39% identity with the branching enzyme from *Thermus thermophilus*, which is the only GlgB belonging to the GH57 family in CAZy [54] for which both kinetic and structural data are available [55]. Curiously, the putative GlgB from *Melainabacteria* showed 93% identity with a putative homologous enzyme from *Clostridium* (see supplemental Figure S3), but only 30 and 40% identity with GH57 GlgBs from *Bacillus halodurans* and *Thermococcus kodakarensis*, respectively. On the other hand, when the GlgB from *Anabaena* (belonging to the CAZy GH13 family, as most of the GlgBs already characterized) was used as a template, no significant coincidences were found in the *Melainabacteria* genomic information. Recently, it was suggested that the GH57 GlgB produces glucans with short branches, although remaining work should be completed to understand the precise role of this type of enzyme [56]. Then, given the presence of genes encoding the complete classical pathway for glycogen synthesis, it can be suggested that the glucan would act as a molecule for carbon and energy storage in *Melainabacteria* [4], possibly with some structural particularities yet to be elucidated [57].

The hypothesis of glycogen as a carbon/energy allocation molecule in *Melainabacteria* is reinforced by the presence of the gene encoding the maltosyl-transferase GlgE (EC 2.4.99.16), an enzyme that elongates a linear α-1,4-glucan in two glucose units [58]. The latter was only characterized from actinobacterial sources, being crystallized the one from *Streptomyces* [59] or proposed as an anti-tuberculosis drug [60]. The presence of genes for two glycogen pathways was postulated after *in silico* analysis [61], demonstrated in *M. tuberculosis* [62,63], and very recently biochemically characterized in *Chlamydia* [64]. Also, it was established that the mycobacterial GlgA enzyme catalyzes the synthesis of maltose-1P, the specific substrate for glycogen elongation by GlgE [65]. The substrates for maltose-1P synthesis by the mycobacterial GlgA (now GlgM) are Glc-1P (acceptor) and ADP-Glc (glucosyl donor), both closely linked to ADP-GlcPPase activity, thus strengthening the leading role of the latter in bacterial glycogen metabolism. Altogether, the biochemical characterization of ADP-GlcPPases from *Melainabacteria* and the analysis of genes co-existing in their genome allowed us to postulate that carbon management in these bacteria is similar to that from other heterotrophic microorganisms, particularly Actinobacteria. On the other hand, we found no genes related to the synthesis of sucrose in *Melainabacteria* [*e*.*g*., sucrose-6P synthase (EC 2.4.1.14) and sucrose synthase (EC 2.4.1.13)], as it occurs in *Anabaena* (see Table 2). Thus, we hypothesize that this might be a critical difference between *Anabaena* (and other cyanobacterial organisms) with the sister-clade *Melainabacteria*. This difference might be reflected in the different regulatory properties of the respective ADP-GlcPPases.

Overall, the comparative analysis between ADP-GlcPPases from Anabaena PCC 7120 and *Melainabacteria* would help to discover new allosteric regulators in future biochemical studies. This work emphasizes the importance of understanding the link between the synthesis of storage compounds, like glycogen, with metabolites that indicate the carbon and energy status of the cell, by studying the kinetic, regulatory, and structural features of important metabolic enzymes. To the best of our knowledge, this is the first biochemical report on enzymes involved in metabolic pathways from *Melainabacteria*, adding data to the hot topic related to the separation of photosynthetic and non-photosynthetic cyanobacteria.

## Acknowledgments

This work was supported by grants from ANPCyT (PICT’17 1515 and PICT’18 00929 to AAI; PICT’18 00698 to MDAD and PICT’18 00865 to CMF), National Science Foundation (grants MCB 1616851 to MAB) and CONICET (PUE-2016-0040 to IAL). MVF is a Fellow from CONICET. RAH was a recipient of a Scholarship from Al Baha University, Saudi Arabia. CMF, AAI and MDAD are Career Investigator members from CONICET.

## Contribution

CMF, MAB, AAI and MDAD designed the work; MVF, RH, MDAD data collection; CMF, MAB, AAI and MDAD analyzed the data; MVF, CMF, MAB, AAI and MDAD wrote the manuscript; all authors have approved the final article.

