## Supplemental Material for "The ADP-glucose pyrophosphorylase from *Melainabacteria*: a comparative study between photosynthetic and non-photosynthetic bacterial sources"

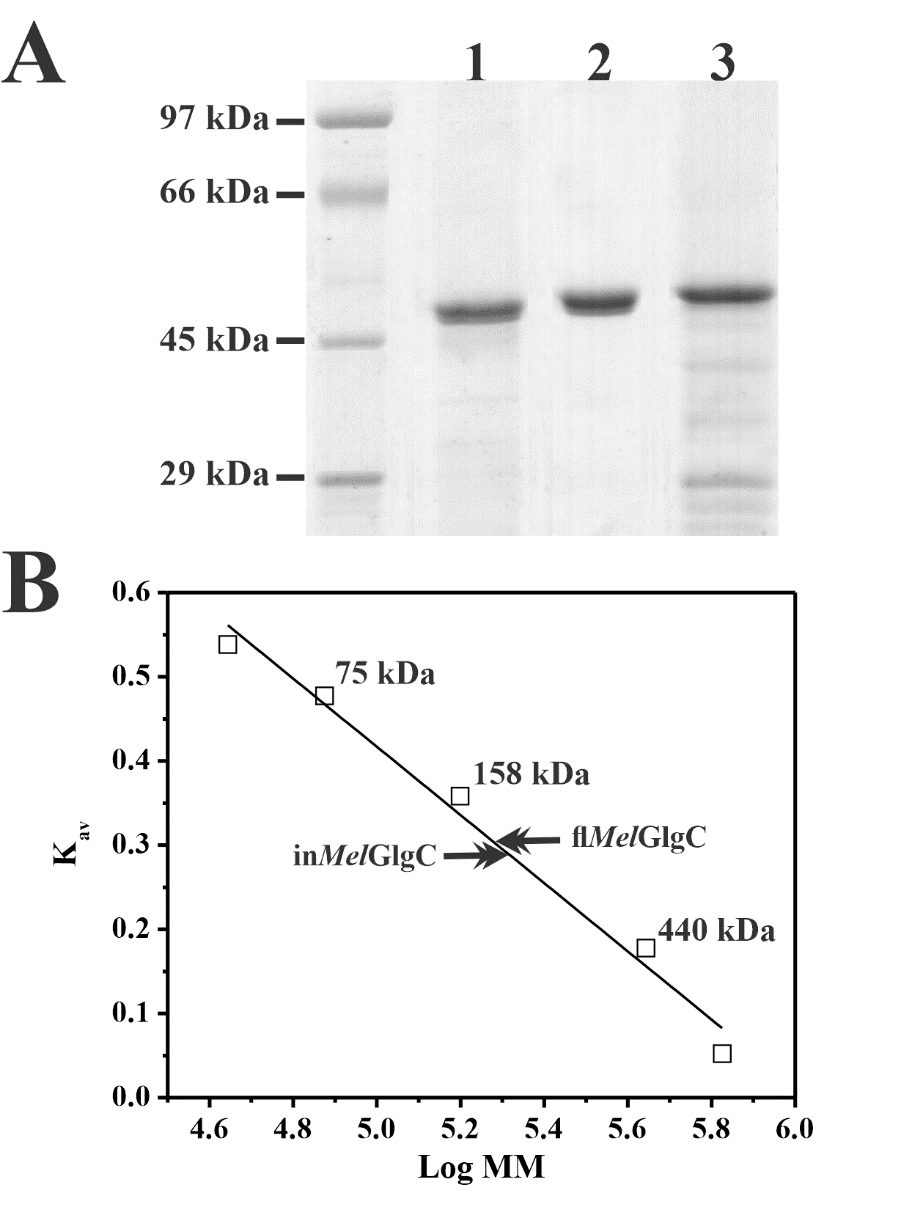

**Supplemental Figure S1. Structural analysis of recombinant ADP-GlcPPases**. **(A)** SDS-PAGE of purified recombinant ADP-GlcPPases. Lane 1: molecular mass markers; lane 2: His-tagged in*Mel*GlgC; lane 3: His-tagged fl*Mel*GlgC; lane 4: His-tagged *Ana*GlgC. **(B)** Molecular mass (MM) determination of the purified proteins by size exclusion chromatography on Superdex 200.

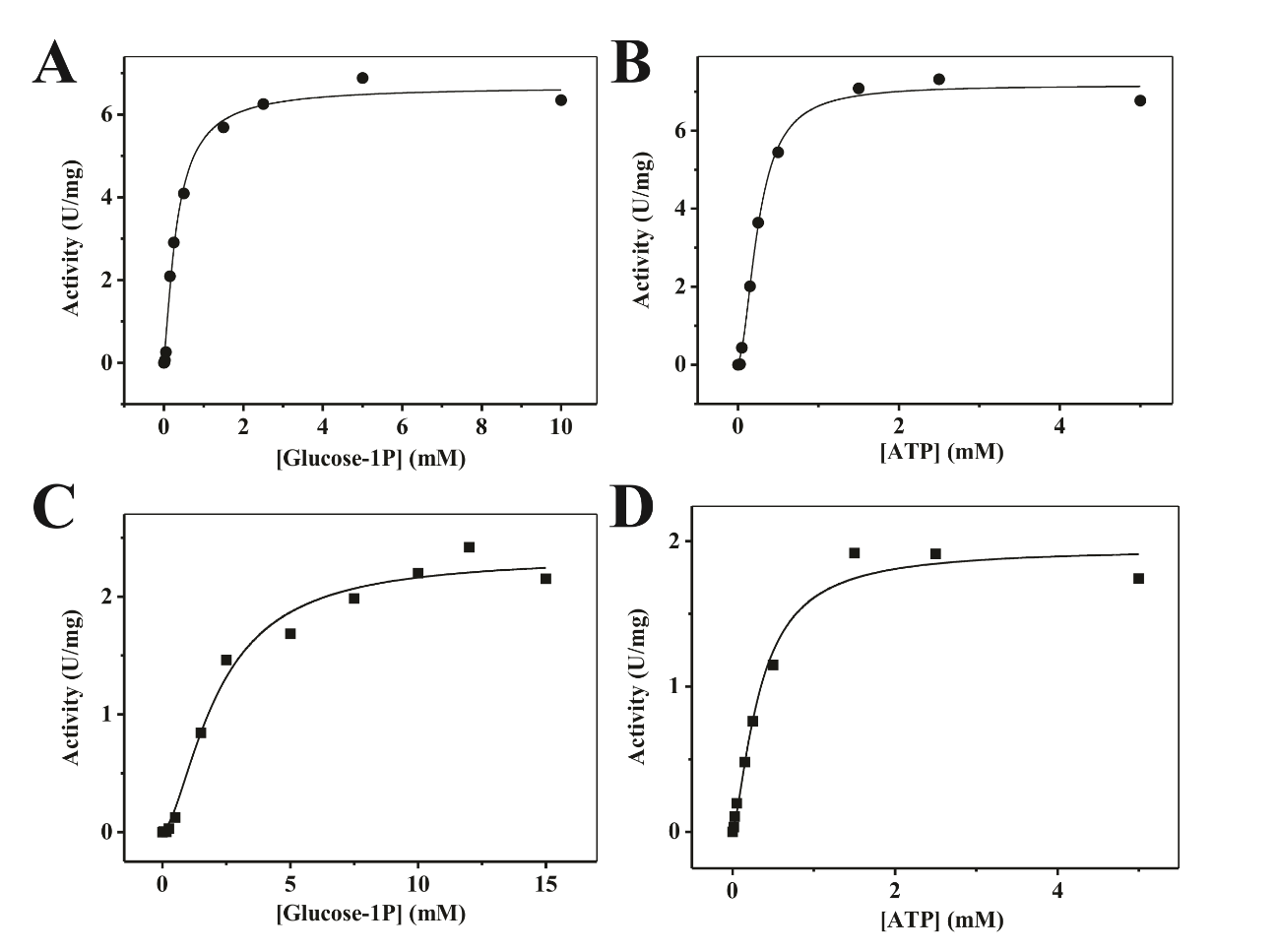

**Supplemental Figure S2:** Glc-1P and ATP saturation curves for intestinal (A-B, filled circles) and free-living (C-D, filled squares) *Melainabacteria* ADP-GlcPPases.

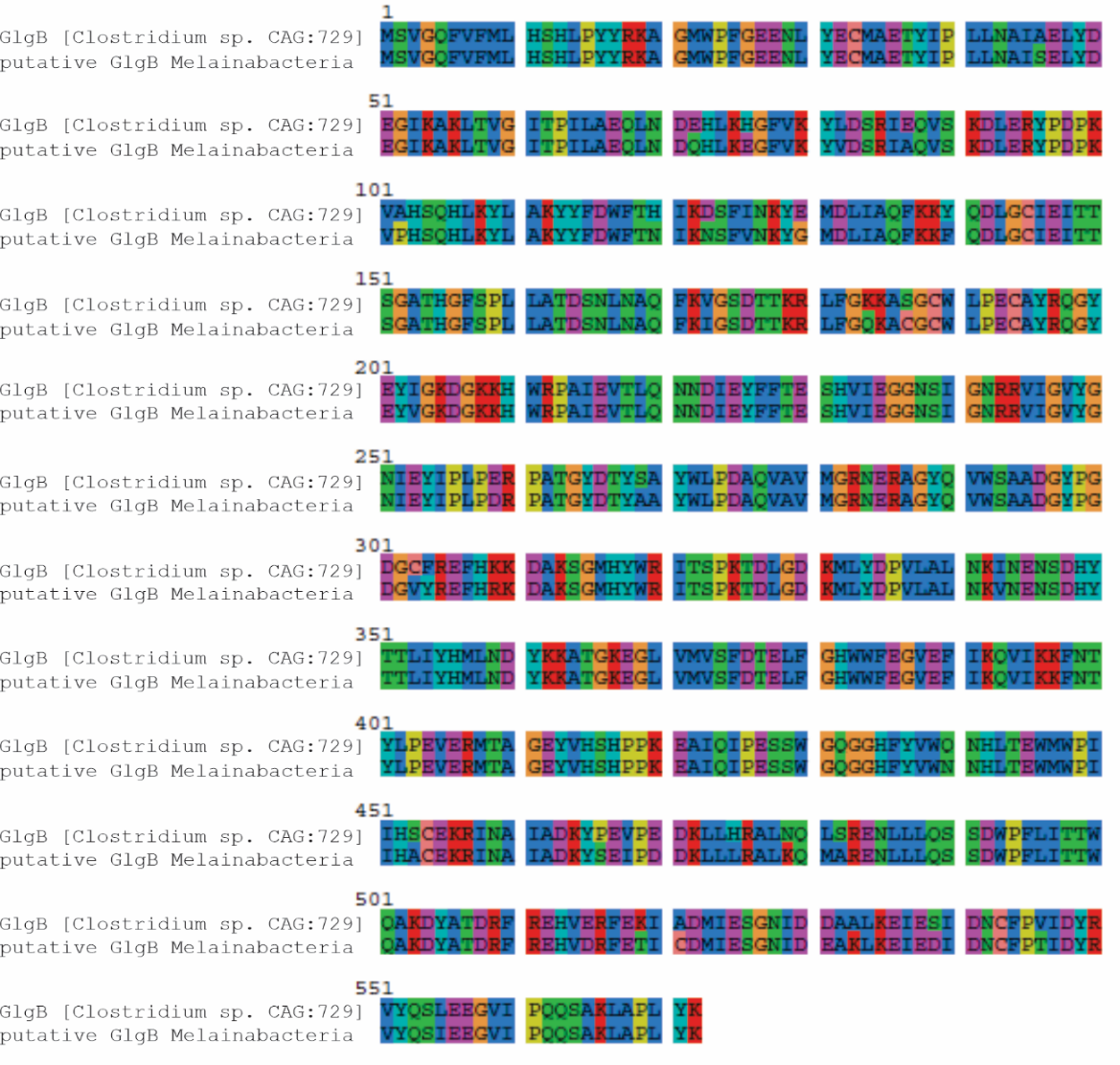

**Supplemental Figure S3**: Sequence alignment of GlgB from *Clostridium* with the putative GlgB from *Melainabacteria*.

**Supplemental Table S1:** Kinetic parameters for allosteric effectors of cyanobacterial ADP-GlcPPases.

| **Effector** | **in*Mel*GlgC** | | **fl*Mel*GlgC** | | ***Ana*GlgC** | |
| --- | --- | --- | --- | --- | --- | --- |
|  | *A*_0.5_ (mM) | Activation^a^  (-fold) | *A*_0.5_ (mM) | Activation^a^  (-fold) | *A*_0.5_ (mM) | Activation^a^  (-fold) |
| **Glc-6P** | 0.37 ± 0.01 | 54 | 0.35 ± 0.03 | 11.7 | 1.45 ± 0.16 | 5.4 |
| **Fru-6P** | 0.44 ± 0.05 | 40 | 1.7 ± 0.1 | 3.8 | 0.9 ± 0.1 | 14.4 |
| **Man-6P** | 0.52 ± 0.02 | 14.7 | 0.45 ± 0.04 | 12.8 | 0.40 ± 0.06 | 6 |
| **3-PGA** | *No effect* | 1 | *No effect* | 1 | 0.19 ± 0.09 | 28.7 |

^a^Activation was calculated as the ratio of *V*_max_ determined from the effector saturation curve over the *V*_max_ of the enzyme in the absence of effector.

**Supplemental Table S2:** Kinetic parameters for inhibitors of cyanobacterial ADP-GlcPPases.

| **Effector** | **in*Mel*GlgC** | | **fl*Mel*GlgC** | | ***Ana*GlgC** | |
| --- | --- | --- | --- | --- | --- | --- |
|  | *I*_0.5_ (mM) | *I*_0.5_ (with 0.4 mM Glc-6P) | *I*_0.5_ (mM) | *I*_0.5_ (with 0.4 mM Glc-6P) | *I*_0.5_ (mM) | *I*_0.5_ (with 0.2 mM 3-PGA) |
| **Pi** | 0.23 ± 0.10 | 0.17 ± 0.06 | 2.3 ± 0.7 | 0.8 ± 0.3 | 0.090 ± 0.004 | 0.20 ± 0.07 |
| **ADP** | 2.3 ± 0.9 | 1.3 ± 0.2 | 1.44 ± 0.08 | 2.7 ± 0.3 | 1.07 ± 0.09 | 0.9 ± 0.1 |

| **Supplemental Table S3:** Number codes for sequences used to build the phylogenetic tree (showed in Figure 1). Color highlighting links numbers with that showen in the tree representing different taxonomies | |
| --- | --- |
| 1 | gi\|62738704\|pdb\|1YP2\|A Chain A, Crystal Structure Of Potato Tuber Adp-Glucose Pyrophosphorylase [Solanum tuberosum] |
| 2 | gi\|15238933\|ref\|NP_199641.1\| glucose-1-phosphate adenylyltransferase small subunit [Arabidopsis thaliana] |
| 3 | gi\|357495273\|ref\|XP_003617925.1\| Glucose-1-phosphate adenylyltransferase [Medicago truncatula] |
| 4 | gi\|356552274\|ref\|XP_003544493.1\| PREDICTED: glucose-1-phosphate adenylyltransferase small subunit, chloroplastic-like isoform 1 [Glycine max] |
| 5 | gi\|224131934\|ref\|XP_002321214.1\| predicted protein [Populus trichocarpa] |
| 6 | gi\|255567204\|ref\|XP_002524583.1\| glucose-1-phosphate adenylyltransferase, putative [Ricinus communis] |
| 7 | gi\|225447450\|ref\|XP_002263255.1\| PREDICTED: hypothetical protein [Vitis vinifera] |
| 8 | gi\|556622\|emb\|CAA55515.1\| ADP-glucose pyrophosphorylase [Beta vulgaris subsp. vulgaris] |
| 9 | gi\|1237080\|emb\|CAA65539.1\| ADP-glucose pyrophosphorylase [Pisum sativum] |
| 10 | gi\|440595\|emb\|CAA54260.1\| ADP-glucose pyrophosphorylase [Vicia faba var. minor] |
| 11 | gi\|162462257\|ref\|NP_001105178.1\| ADP-glucose pyrophosphorylase small subunit [Zea mays] |
| 12 | gi\|115476014\|ref\|NP_001061603.1\| Os08g0345800 [Oryza sativa Japonica Group] |
| 13 | gi\|357157910\|ref\|XP_003577955.1\| PREDICTED: glucose-1-phosphate adenylyltransferase small subunit [Brachypodium distachyon] |
| 14 | gi\|242048788\|ref\|XP_002462140.1\| hypothetical protein SORBIDRAFT_02g020410 [Sorghum bicolor] |
| 15 | gi\|27464770\|gb\|AAO16183.1\| ADP-glucose pyrophosphorylase small subunit [Hordeum vulgare subsp. vulgare] |
| 16 | gi\|52430025\|gb\|AAU50665.1\| ADP-glucose pyrophosphorylase small subunit [Triticum aestivum] |
| 17 | gi\|232166\|sp\|Q00081.1\|GLGL1_SOLTU RecName: Full=Glucose-1-phosphate adenylyltransferase large subunit 1 [Solanum tuberosum] |
| 18 | gi\|14916987\|sp\|P55229.3\|GLGL1_ARATH RecName: Full=Glucose-1-phosphate adenylyltransferase large subunit 1 [Arabidopsis thaliana] |
| 19 | gi\|1947084\|gb\|AAC49941.1\| ADP-glucose pyrophosphorylase large subunit agpl1 [Solanum lycopersicum] |
| 20 | gi\|297812109\|ref\|XP_002873938.1\| hypothetical protein ARALYDRAFT_488807 agpl1 [Arabidopsis lyrata subsp. lyrata] |
| 21 | gi\|297821353\|ref\|XP_002878559.1\| predicted protein agpl4 [Arabidopsis lyrata subsp. lyrata] |
| 22 | gi\|357467317\|ref\|XP_003603943.1\| Glucose-1-phosphate adenylyltransferase large subunit [Medicago truncatula] |
| 23 | gi\|356571037\|ref\|XP_003553688.1\| PREDICTED: glucose-1-phosphate adenylyltransferase large subunit 2, chloroplastic-like [Glycine max] |
| 24 | gi\|356563435\|ref\|XP_003549968.1\| PREDICTED: glucose-1-phosphate adenylyltransferase large subunit 1, chloroplastic-like [Glycine max] |
| 25 | gi\|356562361\|ref\|XP_003549440.1\| PREDICTED: glucose-1-phosphate adenylyltransferase large subunit, chloroplastic/amyloplastic-like [Glycine max] |
| 26 | gi\|356517038\|ref\|XP_003527197.1\| PREDICTED: glucose-1-phosphate adenylyltransferase large subunit 1-like [Glycine max] |
| 27 | gi\|162460455\|ref\|NP_001106017.1\| plastid ADP-glucose pyrophosphorylase large subunit [Zea mays] |
| 28 | gi\|189027076\|ref\|NP_001121104.1\| glucose-1-phosphate adenylyltransferase large subunit 1, chloroplastic/amyloplastic [Zea mays] |
| 29 | gi\|357132398\|ref\|XP_003567817.1\| PREDICTED: glucose-1-phosphate adenylyltransferase large subunit [Brachypodium distachyon] |
| 30 | gi\|115455167\|ref\|NP_001051184.1\| Os03g0735000 [Oryza sativa Japonica Group] |
| 31 | gi\|115471355\|ref\|NP_001059276.1\| Os07g0243200 [Oryza sativa Japonica Group] |
| 32 | gi\|242088961\|ref\|XP_002440313.1\| hypothetical protein SORBIDRAFT_09g029610 [Sorghum bicolor] |
| 33 | gi\|2105137\|gb\|AAC49729.1\| ADP-glucose pyrophosphorylase large subunit [Hordeum vulgare subsp. vulgare] |
| 34 | gi\|22347636\|gb\|AAM95945.1\| ADP-glucose pyrophosphorylase large subunit [Oncidium Goldiana] |
| 35 | gi\|32812836\|emb\|CAD98749.1\| ADP-glucose pyrophosphorylase large subunit [Triticum aestivum] |
| 36 | gi\|1707930\|sp\|P12299.2\|GLGL2_WHEAT RecName: Full=Glucose-1-phosphate adenylyltransferase large subunit [Triticum aestivum] |
| 37 | gi\|87124328\|ref\|ZP_01080177.1\| ADP-glucose pyrophosphorylase [Synechococcus sp. RS9917] |
| 38 | gi\|126660345\|ref\|ZP_01731458.1\| glucose-1-phosphate adenylyltransferase [Cyanothece sp. CCY0110] |
| 39 | gi\|318041355\|ref\|ZP_07973311.1\| glucose-1-phosphate adenylyltransferase [Synechococcus sp. CB0101] |
| 40 | gi\|22298830\|ref\|NP_682077.1\| glucose-1-phosphate adenylyltransferase [Thermosynechococcus elongatus BP-1] |
| 41 | gi\|186686123\|ref\|YP_001869319.1\| glucose-1-phosphate adenylyltransferase [Nostoc punctiforme PCC 73102] |
| 42 | gi\|17232137\|ref\|NP_488685.1\| glucose-1-phosphate adenylyltransferase [Nostoc sp. PCC 7120] |
| 43 | gi\|16332282\|ref\|NP_443010.1\| glucose-1-phosphate adenylyltransferase [Synechocystis sp. PCC 6803] |
| 44 | gi\|56750930\|ref\|YP_171631.1\| glucose-1-phosphate adenylyltransferase [Synechococcus elongatus PCC 6301] |
| 45 | gi\|75908241\|ref\|YP_322537.1\| glucose-1-phosphate adenylyltransferase [Anabaena variabilis ATCC 29413] |
| 46 | gi\|170076729\|ref\|YP_001733367.1\| glucose-1-phosphate adenylyltransferase [Synechococcus sp. PCC 7002] |
| 47 | gi\|307151922\|ref\|YP_003887306.1\| glucose-1-phosphate adenylyltransferase [Cyanothece sp. PCC 7822] |
| 48 | gi\|298492804\|ref\|YP_003722981.1\| glucose-1-phosphate adenylyltransferase ['Nostoc azollae' 0708] |
| 49 | gi\|209527099\|ref\|ZP_03275613.1\| glucose-1-phosphate adenylyltransferase [Arthrospira maxima CS-328] |
| 50 | gi\|303273364\|ref\|XP_003056043.1\| adp-glucose pyrophosphorylase [Micromonas pusilla CCMP1545] |
| 51 | gi\|303271247\|ref\|XP_003054985.1\| adp-glucose pyrophosphorylase [Micromonas pusilla CCMP1545] |
| 52 | gi\|255070935\|ref\|XP_002507549.1\| adp-glucose pyrophosphorylase [Micromonas sp. RCC299] |
| 53 | gi\|255080070\|ref\|XP_002503615.1\| glucose-1-phosphate adenylyltransferase [Micromonas sp. RCC299] |
| 54 | gi\|159470605\|ref\|XP_001693447.1\| ADP-glucose pyrophosphorylase large subunit [Chlamydomonas reinhardtii] |
| 55 | gi\|159467349\|ref\|XP_001691854.1\| ADP-glucose pyrophosphorylase small subunit [Chlamydomonas reinhardtii] |
| 56 | gi\|308814250\|ref\|XP_003084430.1\| AGPLU2 (ISS) [Ostreococcus tauri] |
| 57 | gi\|308806175\|ref\|XP_003080399.1\| AGPSU1 (ISS) [Ostreococcus tauri] |
| 58 | gi\|302849075\|ref\|XP_002956068.1\| hypothetical protein VOLCADRAFT_76956 [Volvox carteri f. nagariensis] |
| 59 | gi\|302840808\|ref\|XP_002951950.1\| hypothetical protein VOLCADRAFT_75183 [Volvox carteri f. nagariensis] |
| 60 | gi\|145356323\|ref\|XP_001422382.1\| predicted protein [Ostreococcus lucimarinus CCE9901] |
| 61 | gi\|145349062\|ref\|XP_001418959.1\| predicted protein [Ostreococcus lucimarinus CCE9901] |
| 62 | gi\|302825850\|ref\|XP_002994500.1\| hypothetical protein SELMODRAFT_138695 [Selaginella moellendorffii] |
| 63 | gi\|302815217\|ref\|XP_002989290.1\| hypothetical protein SELMODRAFT_129625 [Selaginella moellendorffii] |
| 64 | gi\|302802313\|ref\|XP_002982912.1\| hypothetical protein SELMODRAFT_117069 [Selaginella moellendorffii] |
| 65 | gi\|302800351\|ref\|XP_002981933.1\| hypothetical protein SELMODRAFT_115472 [Selaginella moellendorffii] |
| 66 | gi\|302798196\|ref\|XP_002980858.1\| hypothetical protein SELMODRAFT_233627 [Selaginella moellendorffii] |
| 67 | gi\|302788037\|ref\|XP_002975788.1\| hypothetical protein SELMODRAFT_267891 [Selaginella moellendorffii] |
| 68 | gi\|302783933\|ref\|XP_002973739.1\| hypothetical protein SELMODRAFT_149205 [Selaginella moellendorffii] |
| 69 | gi\|302773934\|ref\|XP_002970384.1\| hypothetical protein SELMODRAFT_231637 [Selaginella moellendorffii] |
| 70 | gi\|302769466\|ref\|XP_002968152.1\| hypothetical protein SELMODRAFT_169778 [Selaginella moellendorffii] |
| 71 | WP_119256651.1 glucose-1-phosphate adenylyltransferase [Shinella zoogloeoides] |
| 72 | WP_115756538.1 glucose-1-phosphate adenylyltransferase [Paracoccus thiocyanatus] |
| 73 | WP_115156201.1 glucose-1-phosphate adenylyltransferase [Rhizobiales bacterium] |
| 74 | WP_114579936.1 glucose-1-phosphate adenylyltransferase [Mesorhizobium oceanicum] |
| 75 | WP_111356150.1 glucose-1-phosphate adenylyltransferase [Rhodoplanes elegans] |
| 76 | WP_109150954.1 glucose-1-phosphate adenylyltransferase [Azospirillum sp. TSO5] |
| 77 | WP_106310289.1 glucose-1-phosphate adenylyltransferase [Martelella mediterranea] |
| 78 | WP_028746729.1 glucose-1-phosphate adenylyltransferase [Rhizobium mesoamericanum] |
| 79 | WP_014493115.1 glucose-1-phosphate adenylyltransferase [Bradyrhizobium japonicum] |
| 80 | WP_113529535.1 glucose-1-phosphate adenylyltransferase [Rhizobiales bacterium] |
| 81 | WP_126462236.1 glucose-1-phosphate adenylyltransferase [Sulfuritortus calidifontis] |
| 82 | WP_090633633.1 glucose-1-phosphate adenylyltransferase [Nitrosomonas marina] |
| 83 | WP_090317903.1 glucose-1-phosphate adenylyltransferase [Nitrosomonas oligotropha] |
| 84 | WP_110464968.1 glucose-1-phosphate adenylyltransferase [Xylophilus ampelinus] |
| 85 | WP_102635427.1 glucose-1-phosphate adenylyltransferase [Paraburkholderia rhynchosiae] |
| 86 | WP_089164405.1 glucose-1-phosphate adenylyltransferase [Caballeronia sordidicola] |
| 87 | WP_075585858.1 glucose-1-phosphate adenylyltransferase [Rhodoferax antarcticus] |
| 88 | WP_113067136.1 glucose-1-phosphate adenylyltransferase [Nitrosospira multiformis] |
| 89 | WP_096777558.1 glucose-1-phosphate adenylyltransferase [Neisseria dumasiana] |
| 90 | WP_111935803.1 glucose-1-phosphate adenylyltransferase [Paraburkholderia bryophila] |
| 91 | WP_120645735.1 glucose-1-phosphate adenylyltransferase [Corallococcus sp. CA051B] |
| 92 | WP_028321096.1 glucose-1-phosphate adenylyltransferase [Desulfatiglans anilini] |
| 93 | WP_015352759.1 glucose-1-phosphate adenylyltransferase [Myxococcus stipitatus] |
| 94 | WP_013935338.1 glucose-1-phosphate adenylyltransferase [Myxococcus macrosporus] |
| 95 | WP_002610372.1 glucose-1-phosphate adenylyltransferase [Stigmatella aurantiaca] |
| 96 | WP_050724205.1 glucose-1-phosphate adenylyltransferase [Vulgatibacter incomptus] |
| 97 | WP_013376533.1 glucose-1-phosphate adenylyltransferase [Stigmatella aurantiaca] |
| 98 | WP_012529339.1 glucose-1-phosphate adenylyltransferase [Geobacter bemidjiensis] |
| 99 | WP_124941924.1 glucose-1-phosphate adenylyltransferase [Vibrio mediterranei] |
| 100 | WP_122029635.1 glucose-1-phosphate adenylyltransferase [Salmonella enterica] |
| 101 | WP_119560420.1 glucose-1-phosphate adenylyltransferase [Vibrio cholerae] |
| 102 | WP_118873727.1 glucose-1-phosphate adenylyltransferase [Haemophilus influenzae] |
| 103 | WP_118861127.1 glucose-1-phosphate adenylyltransferase [Haemophilus haemolyticus] |
| 104 | WP_118836321.1 glucose-1-phosphate adenylyltransferase [Aeromonas hydrophila] |
| 105 | WP_117373127.1 glucose-1-phosphate adenylyltransferase [Klebsiella pneumoniae] |
| 106 | WP_115431588.1 glucose-1-phosphate adenylyltransferase [Escherichia coli] |
| 107 | WP_016678934.1 glucose-1-phosphate adenylyltransferase, partial [Yersinia pestis] |
| 108 | WP_032403021.1 glucose-1-phosphate adenylyltransferase [Rhodococcus fascians] |
| 109 | WP_073370619.1 glucose-1-phosphate adenylyltransferase [Rhodococcus jostii] |
| 110 | NP_625258.2 glucose-1-phosphate adenylyltransferase [Streptomyces coelicolor A3(2)] |
| 111 | WP_037698817.1 glucose-1-phosphate adenylyltransferase [Streptomyces scabiei] |
| 112 | WP_039674867.1 glucose-1-phosphate adenylyltransferase [Corynebacterium minutissimum] |
| 113 | WP_058598057.1 glucose-1-phosphate adenylyltransferase [Microbacterium testaceum] |
| 114 | WP_045280110.1 glucose-1-phosphate adenylyltransferase [Microbacterium oxydans] |
| 115 | WP_085292363.1 glucose-1-phosphate adenylyltransferase [Mycobacterium vulneris] |
| 116 | WP_096441957.1 glucose-1-phosphate adenylyltransferase [Mycobacterium shigaense] |
| 117 | WP_057375489.1 glucose-1-phosphate adenylyltransferase [Mycobacterium tuberculosis] |
| 118 | WP_036383376.1 glucose-1-phosphate adenylyltransferase [Micrococcus luteus] |
| 119 | WP_092537108.1 glucose-1-phosphate adenylyltransferase [Actinomyces ruminicola] |
| 120 | WP_060958022.1 glucose-1-phosphate adenylyltransferase [Actinomyces oris] |
| 121 | WP_058872255.1 glucose-1-phosphate adenylyltransferase [Kocuria palustris] |
| 122 | >tr\|A0A327JBB5\|Glucose-1-phosphate adenylyltransferase [Candidatus Melainabacteria bacterium] |
| 123 | >tr\|A0A327J7S7\|Glucose-1-phosphate adenylyltransferase [Candidatus Melainabacteria bacterium] |
| 124 | >tr\|A0A327J5Y0\|Glucose-1-phosphate adenylyltransferase [Candidatus Melainabacteria bacterium] |
| 125 | >tr\|A0A292SQB6\|Glucose-1-phosphate adenylyltransferase [Candidatus Gastranaerophilales bacterium HUM_20] |
| 126 | >tr\|A0A1D7YQL5\|Glucose-1-phosphate adenylyltransferase [Candidatus Melainabacteria bacterium MEL.A1] |
| 127 | >tr\|A0A316NNB5\|Glucose-1-phosphate adenylyltransferase [Candidatus Gastranaerophilales bacterium] |
| 128 | >tr\|A0A292SN60\|Glucose-1-phosphate adenylyltransferase [Candidatus Gastranaerophilales bacterium HUM_21] |
| 129 | >tr\|A0A292RFB6\|Glucose-1-phosphate adenylyltransferase [Candidatus Gastranaerophilales bacterium HUM_9] |
| 130 | >tr\|A0A292R804\|Glucose-1-phosphate adenylyltransferase [Candidatus Gastranaerophilales bacterium HUM_7] |
| 131 | >tr\|A0A292R3M3\|Glucose-1-phosphate adenylyltransferase [Candidatus Gastranaerophilales bacterium HUM_4] |
| 132 | >tr\|A0A292QNQ1\|Glucose-1-phosphate adenylyltransferase [Candidatus Gastranaerophilales bacterium HUM_1] |
| 133 | >tr\|A0A1F6PWP2\|Glucose-1-phosphate adenylyltransferase [Candidatus Melainabacteria bacterium GWF2_37_15] |
